# Effects of High Fat Diet on Metabolic Health Vary by Age of Menopause Onset

**DOI:** 10.1101/2024.01.18.576269

**Authors:** Abigail E. Salinero, Harini Venkataganesh, Charly Abi-Ghanem, David Riccio, Richard D. Kelly, Olivia J. Gannon, Avi Sura, Heddwen L. Brooks, Kristen L. Zuloaga

## Abstract

Menopause accelerates metabolic dysfunction, including (pre-)diabetes, obesity and visceral adiposity. However, the effects of endocrine vs. chronological aging in this progression are poorly understood. We hypothesize that menopause, especially in the context of middle-age, will exacerbate the metabolic effects of a high fat diet. Using young-adult and middle-aged C57BL/6J female mice, we modeled diet-induce obesity via chronic administration of high fat (HF) diet vs. control diet. We modeled peri-menopause/menopause via injections of 4-vinylcyclohexene diepoxide, which accelerates ovarian failure vs. vehicle. We performed glucose tolerance tests 2.5 and 7 months after diet onset, during the peri-menopausal and menopausal phases, respectively. Peri-menopause increased the severity of glucose intolerance and weight gain in middle-aged, HF-fed mice. Menopause increased weight gain in all mice regardless of age and diet, while chronological aging drove changes in adipose tissue distribution towards more visceral vs. subcutaneous adiposity. These data are in line with clinical data showing that post-menopausal women are more susceptible to metabolic dysfunction and suggest that greater chorological age exacerbates the effects of endocrine aging (menopause). This work highlights the importance of considering both chronological and endocrine aging in studies of metabolic health.

## INTRODUCTION

Menopause is a major endocrinological shift that leaves women vulnerable to the metabolic risk factors for cardiovascular disease [1], a leading cause of death in developed countries. Postmenopausal women are 12% more likely to meet criteria for metabolic syndrome, than premenopausal women [2]. Menopausal transition is associated with weight gain and accumulation of visceral adipose tissue (VAT) in the abdomen [3]. VAT is associated with increased risk for cardiovascular events such as stroke, even in postmenopausal women with a normal body mass index [4].

Menopause is preceded by peri-menopause, which is characterized by menstrual cycle irregularity and large hormone fluctuations [5]. Importantly, peri-menopause also increases body weight and body fat mass [3]. Clinical studies show that deficits in brain glucose metabolism begin in peri-menopause [6]. Unfortunately, our understanding of peri-menopause is largely relegated to observational clinical studies, as rodent menopause studies often utilize an ovariectomy model that induces sudden drops in estrogen and completely bypasses peri-menopause. A model of peri-menopause and menopause can be achieved through administration of 4-vinylcyclohexene diepoxide (VCD), which causes accelerated ovarian failure while leaving ovaries intact [7]. This model includes a preceding phase similar to human peri-menopause, characterized by estrous cycle irregularity, vasomotor symptoms and large fluctuations in estrogen, and the length of this peri-menopausal period varies by the number of days VCD is administered [7]. Also similar to humans, with this model mice retain ovarian tissue, which continues to secrete androgens into circulation post-menopause [7].

Most clinical studies investigating the effects of menopause on metabolic function are conducted in aged women, making it difficult to differentiate the effects of chronological aging from the effects of endocrine aging (menopause) on metabolic function [8]. Conversely, many rodent studies use exclusively young females. In the current study we sought to identify the unique contributions of endocrine vs chronological age on metabolic health in a mouse model of menopause via accelerated ovarian failure. We show that middle-aged menopausal females fed a high fat (HF) diet suffer more severe metabolic impairment than young HF-fed menopausal females.

## METHODS

All experiments were approved by the Albany Medical College Animal Care and Use Committee and in compliance with the ARRIVE guidelines. Female C57BL/6J mice were purchased at ∼10 and ∼32 weeks of age from Jackson Laboratories (Bar Harbor, ME, USA). Following one week of acclimation, during which time they were provided with Purina Lab Diet 5P76, mice were randomized to treatment groups based on cage ID number and received either 160mg/kg 4-vinylcyclohexene diepoxide (VCD; Sigma #V3630) or vehicle (sesame oil) via i.p. injection for 11 consecutive days. One week after the final injection, mice were placed on either a HF diet (60% fat, 5.21 Kcal/g; D12492, Research Diets, New Brunswick, NJ, USA) or a control diet (Ctrl; 10% fat, 3.82 Kcal/g; D12450B, Research Diets, USA). Mice remained on their respective diets for the duration of the study. After 2.5 months on the diet (peri-menopausal phase for VCD injected mice), mice underwent a glucose tolerance test (GTT) as previously described [9], then began daily vaginal cytology to confirm peri-menopausal status. Vaginal cytology continued until acyclic (constant diestrus for >10 days) [10]. After 7 months on diet (menopausal phase for VCD injected mice), mice underwent a second GTT. At the end of the study, visceral and subcutaneous fat pad wet weights were taken. Data were analyzed via 3-way ANOVA with Prism 10.1 (GraphPad Software, San Diego, CA, USA).

## RESULTS

### Metabolic effects of peri-menopause vary by age of onset

Mice were treated with vehicle or VCD for 11 days (to induce menopause with a long peri-menopausal period) starting at 2 months (young onset) or 8 months of age (middle-age onset) and placed on a HF or control diet (timeline in Figure 1A). Average time to menopause after the last VCD injection was 151.3 ± 4.96 days (data not shown). After 2.5 months on their diets, while VCD injected mice were in a peri-menopausal phase (irregularly cycling), mice underwent a GTT (Figure 1B). HF groups showed impaired glucose tolerance (greater area under the curve during GTT) relative to controls (main effect of diet, p<0.0001). Further, peri-menopausal mice showed impaired glucose tolerance relative to pre-menopausal mice (main effect of peri-menopause, p<0.0001). Post-hoc analysis revealed that middle-aged, HF peri-menopausal mice had the most severely impaired glucose tolerance relative to both their age-matched pre-menopausal HF controls (p=0.0020) and to younger peri-menopausal mice (p=0.0379). There was also an overall main effect of age in increasing glucose intolerance (p=0.0259). These data suggest that for middle-aged mice, pronounced glucose tolerance impairments begin during peri-menopause and are worse in those consuming a high fat diet.

**Figure 1.**
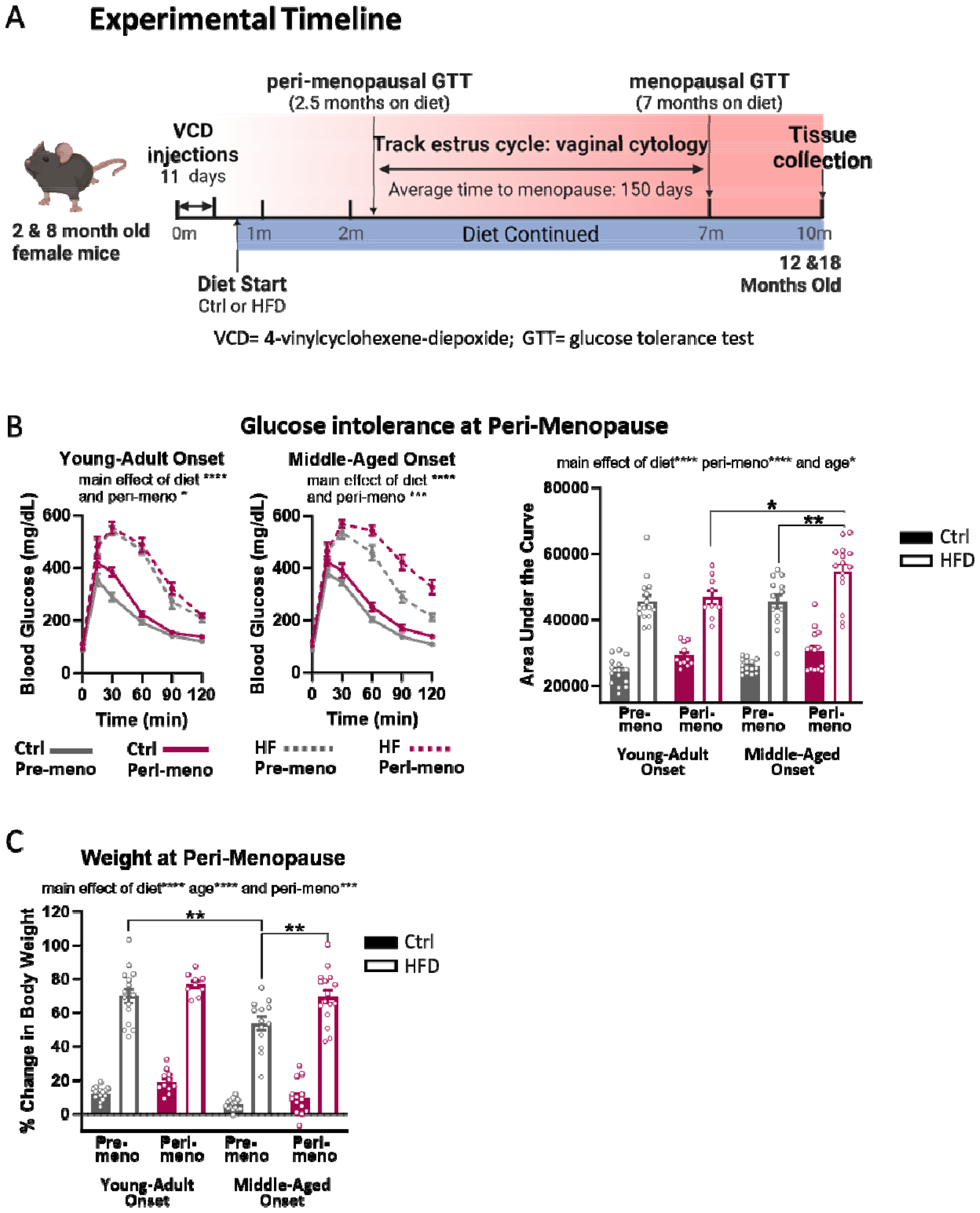
Metabolic effects of peri-menopause vary by age of onset. (A) Experimental Timeline. Two- or 8-month old female C57BL/6J mice were injected for 11 days with VCD or oil. Ten days after final injection, HF or control diet was initiated and continued for the durati**on** of the study. Glucose intolerance was assessed at 2.5 months after diet onset during peri-menopause (irregular cyclicity) via glucose tolerance test (GTT). Vaginal cytology was performed daily to confirm acyclicity. After mice stopped cycling, (by 7 months on diet) the GTT was repeated. Tissues including subcutaneous and visceral adipose tissue were collected when mice were 12- and 18-months old, 10 months after the start of VCD injections. Body weight was measured and % weight gained relative to starting weight was calculated. (B) At 2.5 and 7 months after the onset of the dietary intervention, mice were fasted overnight, and the next morning baseline blood glucose levels in saphenous vein blood were measured by glucometer (Verio IQ, OneTouch, Sunnyvale CA, USA). Each mouse received 2g/kg of glucose via i.p. injection and blood glucose levels were re-measured at 15, 30, 60, 90 and 120 minutes post-injection. Area under the curve (AUC) from the glucose tolerance test was calculated. Blood glucose concentrations (mg/dL) over 2 hours post-glucose challenge for young adult onset (left panel) and middle-aged onset (middle panel) mice. n=9-15/group, repeated measures ANOVA for each age group. (Right panel) AUC measured 2.5 months after diet onset, during “peri-menopausal” period. 3-way ANOVA with Sidak’s multiple comparisons (C) % Change in body weight after 3 months on the HF or Control diet. 3-way ANOVA with Sidak’s multiple comparisons. n=9-15/group, *p<0.05, **p<0.01, ****p<0.0001. Main effects are written above graphs and post-hoc differences appear on graphs. HF = high fat; Ctrl = control; Pre-meno = pre-menopausal; Peri-meno = peri-menopausal; Meno = menopausal

Weight gain during the peri-menopausal phase was measured as a percent change relative to starting weight (Figure 1C). Both HF diet and younger age caused weight gain (main effects of diet and age: p<0.0001). Further, peri-menopausal mice gained more weight than pre-menopausal mice (main effect of peri-menopause: p=0.0002). However, post-hoc tests showed peri-menopausal weight gain was driven most heavily by HF-fed middle-aged onset peri-menopausal mice, which gained significantly more weight than their pre-menopausal counterparts (p=0.0025). These data suggest that, especially at middle-age, differences in weight gain begin at peri-menopause.

### Metabolic effects of menopause vary by age of onset

GTTs were repeated after 7 months on the diet during menopause (once VCD-injected mice were acyclic). GTT area under the curve analysis (Figure 2A) showed that young onset mice—which were middle-aged at the time of this GTT—had more impaired glucose tolerance relative to the older mice (main effect of age, p=0.0461). Further, all HF groups showed impaired glucose tolerance relative to their controls (main effect of diet, p<0.0001). Finally, menopausal mice showed impaired glucose tolerance relative to vehicle-treated mice (main effect of menopause, p=0.0298). These data suggest that menopause impairs glucose tolerance.

**Figure 2.**
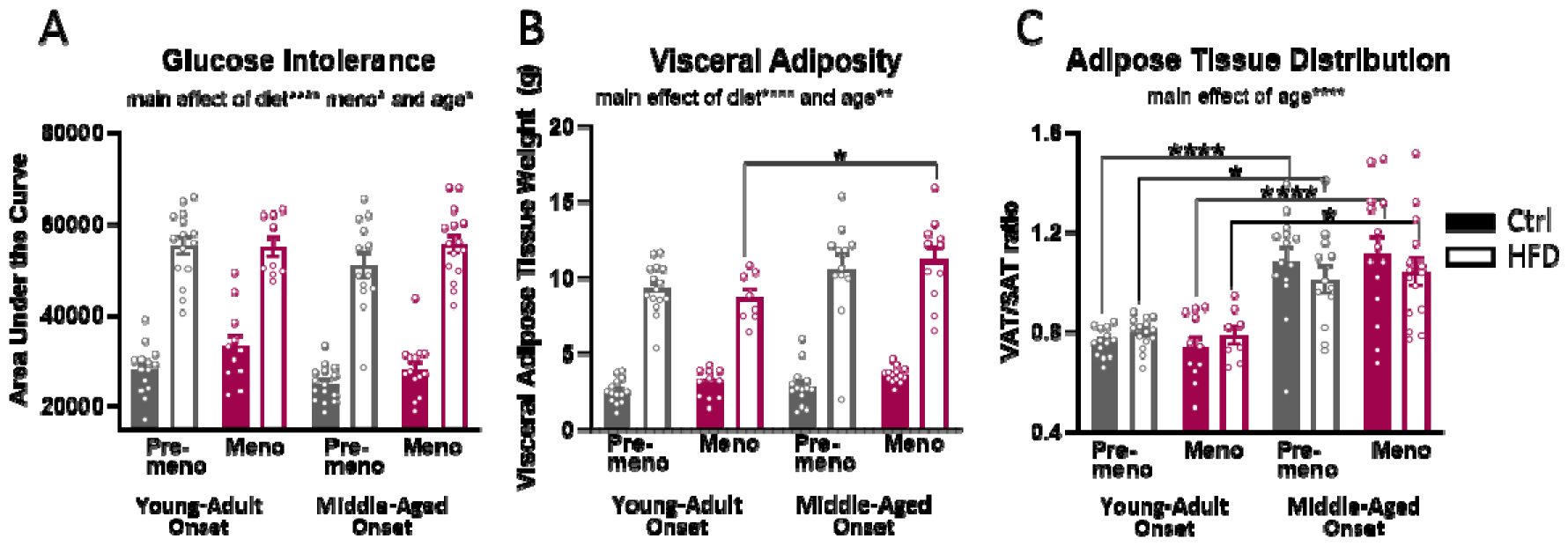
Metabolic effects of menopause vary by age of onset. Glucose tolerance test was repeated after mice had stopped cycling, 7 months after diet onset. (A) AUC performed 7 months after diet onset, during “menopausal” period. (B) Visceral adipose tissue (VAT) wet weights were dissected at the end of the study. (C) Adipose tissue distribution at the end of the study measured as the ratio of visceral vs subcutaneous fat wet weights. n=9-15/group, 3-way ANOVA with Sidak’s post-hoc differences *p<0.05, **p<0.01, ****p<0.0001. Main effects are written above graphs and post-hoc differences appear on graphs. HF = high fat; Ctrl = control; Pre-meno = pre-menopausal; Meno = menopausal; VAT = visceral adipose tissue; SAT = subcutaneous adipose tissue

Adipose tissue was dissected and weighed at the end of the study. Overall, HF diet and age increased visceral adipose tissue (VAT) mass (main effect of diet p<0.0001 and age p=0.0022; Figure 2B). This diet- and age-related increase was strongest in the menopausal mice (HF menopausal middle-aged onset vs HF menopausal young-adult onset p=0.0462). To assess adipose tissue distribution, the ratio of VAT to subcutaneous adipose tissue (SAT) mass was calculated (Figure 2C). Overall, middle-aged onset mice, who were 18 months old at the time of adipose tissue collection, showed a significantly increased VAT:SAT ratio compared to young-adult onset mice, who were 12 months old at the time of tissue collection (main effect of age p<0.0001). Surprisingly, menopausal status did not alter VAT:SAT ratio. Altogether, these data suggest that adipose tissue distribution is driven by chronological age.

## DISCUSSION

Our study shows that when menopause and diet-induced obesity are initiated in young-adult and middle-aged mice, peri-menopause accelerates weight gain and glucose intolerance to the greatest extent in middle-aged, HF-fed mice. However, chronological aging, not menopause, increases the ratio of visceral to subcutaneous fat. Thus, the contributions of chronological and endocrine aging to metabolic health are both distinct and synergistic.

We found that advanced chronological age exacerbates glucose intolerance and weight gain in peri-menopausal mice on a HF diet. Our findings are in agreement with only minimal glucose intolerance effects of peri-menopause in young rodents [11], but increased weight gain following menopause [12, 13]. The current study is the first to directly compare glucose intolerance and weight among young-adult onset and middle-age onset of peri-menopause. In humans, impairments in glucose tolerance often begin with peri-menopause [6], during which large fluctuations in estrogen disrupt estrogen-dependent signaling and transcriptional pathways that regulate energy metabolism and insulin resistance [14]. Our data suggest that middle-age might be a necessary background upon which glucose intolerance is initiated during peri-menopause, thus highlighting the need to use appropriate age in preclinical models.

Some studies indicate that menopausal transition drives increased fat mass and decreased lean mass [15, 16]. Surprisingly, menopause did not alter adipose tissue distribution; instead, chronological age significantly increased VAT:SAT ratio. Notably, in younger females the ratio was on average <1, indicating that there was more SAT than VAT. Among older females the average ratio was >1, indicating they had more VAT. Such age-related body fat redistribution has been described in humans and rodents alike [17]. With age, cellular metabolic function declines in all adipose tissue depots [17]. Subcutaneous adipocytes decrease in number, and the lipid-storing capacity of those that remain is reduced, increasing vulnerability to lipotoxicity. Conversely, VAT is associated with insulin resistance [17]. Our data suggest that this age-related redistribution in favor of visceral adiposity is independent of diet or menopause.

Our current study lays the groundwork for future investigations into the many mechanisms underlying the differences we observed. One possible mechanism for the accelerated metabolic impairments observed with middle-age could be lipotoxicity-induced inflammation and mitochondrial dysfunction. This could be exacerbated with loss of the anti-inflammatory estrogen during the transition to menopause. In support of this, many studies have underlined that a feature common to both chronological aging and menopause is a shift toward chronic, low-grade inflammation [14, 16-18]. Exploration of these mechanisms is critical to the development of efficacious therapeutic strategies to ameliorate metabolic risk factors for a variety of diseases in menopausal women. Our data highlight the importance of considering chronological age in mouse models of menopause.

## DECLARATIONS

### ETHICS APPROVAL AND CONSENT TO PARTICIPATE

Not applicable (no human subjects).

### CONSENT FOR PUBLICATION

Not applicable (no human subjects).

### AVAILABILITY OF DATA AND MATERIALS

The datasets acquired and/or analyzed during the current study are available from the corresponding author on reasonable request.

### COMPETING INTERESTS

The authors have no conflicts to disclose.

### FUNDING

This work was funded by an American Heart Association pre-doctoral award 908878 (AES), BrightFocus Foundation postdoctoral fellowship A2022001F (CAG), NINDS/NIA R01NS110749 (KLZ), NIA U01 AG072464 (KLZ), Alzheimer’s Association AARG-21-849204 (KLZ).

### AUTHORS’ CONTRIBUTIONS

KLZ obtained funding for the experiments. KLZ and AES designed the experiments. AES, CAG, HV, RMS, RDK, OJG performed the animal work. AES analyzed the data. AES prepared the figures. KLZ and AES interpreted the results. AES prepared the manuscript. KLZ edited the manuscript. All authors approved the final manuscript.

## ACKNOWLEDGEMENTS

None.

## Notes

### Competing Interest Statement

The authors have declared no competing interest.

